# Daily Artificial Gravity is Associated with Greater Neural Efficiency during Sensorimotor Adaptation

**DOI:** 10.1101/2022.10.27.514043

**Authors:** G.D. Tays, K.E. Hupfeld, H.R. McGregor, N.E. Gadd, I. S. Kofman, Y. E. De Dios, E.R. Mulder, J.J. Bloomberg, A.P. Mulavara, S.J. Wood, R. D. Seidler

**Affiliations:** Department of Applied Physiology and Kinesiology, University of Florida, Gainesville, FL, USA; KBR, Houston, TX, USA; German Aerospace Center (DLR); NASA Johnson Space Center, Houston, TX, USA; Norman Fixel Institute for Neurological Diseases, University of Florida, Gainesville, FL

**Author notes:** <u>Correspondence to</u>: Rachael Seidler, PhD.

## Abstract

Altered vestibular signaling and body unloading in microgravity results in sensory reweighting and adaptation. Microgravity effects are well-replicated in head-down tilt bed rest (HDBR). Artificial gravity (AG) is a potential countermeasure to mitigate effects of microgravity. We examined the effectiveness of daily AG for mitigating brain and/or behavioral changes in 60 days of HDBR. One group received AG for 30 minutes daily (AG; n=16) and a control group spent the same time in HDBR but received no AG (CTRL; n=8). All participants performed a sensorimotor adaptation task 5 times during fMRI scanning: twice prior to HDBR twice during HDBR, and once following HDBR. The AG group showed similar behavioral adaptation effects compared with the CTRLs. We identified decreased brain activation in the AG group from pre to late HDBR in the cerebellum for the task baseline portion and in the thalamus, calcarine, cuneus, premotor cortices, and superior frontal gyrus in the AG group during the early adaptation phase. The two groups also exhibited differential brain-behavior correlations. Together, these results suggest that AG may result in a reduced recruitment of brain activity for basic motor processes and sensorimotor adaptation. These effects may stem from somatosensory and vestibular stimulation from AG.

## 1. Introduction

In the microgravity, such as aboard the International Space Station (ISS), astronauts must adapt to various challenges, such as the lack of normal gravity, axial body unloading, headward fluid shifts, elevated levels of CO_2_ that can occur in a closed environment, and disrupted circadian rhythms (c.f. Clement et al., 2020). These stressors have been shown to result in various cognitive impairments, such as dual-tasking deficits, decreased spatial awareness and complaints of “space fog” (Clement et al., 1987; Bock et al., 2003; 2010; Strangman et al., 2014; Garret-Bakelman et al., 2019). Sensorimotor control deficits have also been well documented during spaceflight including increased surgical operative time (Campbell et al., 2005) and increased movement duration and movement amplitude (Bock et al., 2003). During spaceflight, astronauts must adapt their movements and learn to perform them in a new environment (such as controlling a robotic arm), so maintaining high performance is paramount. Following spaceflight, astronauts also experience transient deficits in locomotion (Tays et al., 2021; Mulavara et al., 2018; Miller et al., 2018; Layne et al., 1998; McDonald et al., 1996; Bloomberg et al., 1997), balance (Reschke et al., 1994a; 1994b; 1998; Paloski et al., 1992; 1994; Black et al., 1995; 1999) and bimanual coordination (Tays et al., 2021; Miller et al., 2018; Mulavara et al., 2018). These postflight behavioral changes are believed to be a result of central nervous system adaptation to microgravity. Without gravity, vestibular inputs (particularly from the otoliths) indicating head tilts are absent and get down-weighted (Boyle, 2021; Hupfeld et al., 2022). Upon returning to Earth, these adaptive changes are initially maladaptive for movement in Earth’s gravity; performance typically returns to baseline levels within a few weeks postflight (Miller et al., 2018; Mulavara et al., 2010; Tays et al., 2021).

Head-down tilt bedrest (HDBR) is commonly used as an analog to investigate various effects of spaceflight. Participants lay with their head at 6° below their feet for up to several months at a time to simulate the headward fluid shifts and axial unloading of microgravity. HDBR has been shown to model some of the physiological and functional effects of spaceflight (Hargens and Vico, 2016; Reschke et al., 2009; Mulder et al., 2014; Koppelmans et al., 2015, 2017; Miller et al., 2018; Mulavara et al., 2018; Tays et al., 2022). Sensorimotor adaptation occurs in response to sensory prediction errors, i.e. when a movement does not result in the anticipated sensory feedback. This occurs in the microgravity environment, as head tilts do not result in the anticipated vestibular sensory inputs. Testing sensorimotor adaptation abilities during spaceflight and/or HDBR allows study of how modified sensory inputs may affect motor task performance. We have previously investigated the effects of HDBR plus elevated CO_2_ on performance of a sensorimotor adaptation task, in which subjects use a joystick to perform target directed movement with 45° clockwise rotated visual feedback (Banker et al., 2021; Salazar et al., 2021). We found that HDBR+ CO_2_ resulted in a lack of savings, or memory of prior experience with the same perturbation (Banker et al. 2021). In that same study, we observed decreased activation in the right parahippocampal gyrus, right putamen and left hippocampus during early adaptation and increased activation in the right fusiform gyrus and right caudate nucleus during late adaptation (Salazar et al., 2021). Overall, our prior work suggests that HDBR affects sensorimotor adaptation. This may be in part due to multiple adaptations occurring at once, such as the adaptation to vestibular input changes (from HDBR) at the same time as adapting to the rotated visual feedback.

While sensorimotor-specific countermeasures are under development to mitigate the negative impacts of microgravity, an ideal countermeasure would target more than one system. Artificial gravity (AG) has been investigated as once such integrated countermeasure, essentially restoring the effect of Earth’s gravity on the body through generation of force by continuously rotating a short-arm centrifuge at individualized rate. AG applied along the long axis of the body could potentially mitigate orthostatic intolerance and muscle and bone atrophy, load proprioceptive and somatosensory sensors, mitigate cardiovascular deconditioning, as well as stimulate the vestibular system (Linnarsson et al., 2015; Hargens et al., 2013). AG is typically well tolerated by participants, resulting in only a small percentage of sessions ended due to discomfort or sickness following 60 minutes of continuous centrifugation daily (Arya et al., 2007). Delivering AG intermittently (6 bouts of 5 minutes) instead of continuously (1 bout of 30 minutes) reduces orthostatic intolerance more following 5 days of HDBR (Linnarsson et al., 2015). The present investigation is part of the larger Artificial Gravity Bed Rest – European Space Agency (“AGBRESA”) study; where we had previously found that those individuals who received AG had marginally better balance and mobility performance than controls who received no AG, suggesting that AG could be an effective sensorimotor countermeasure, but more than 30 minutes per day may be necessary (Tays et al., 2022).

In the present study, we administered the same visuomotor rotation task as in our previous HDBR studies (Banker et al., 2021; Salazar et al., 2021) where visual feedback is rotated by 45 degrees during the adaptation phases. Our primary aim was to examine whether centrifugal AG applied along the long axis of the body alters sensorimotor adaptation and associated brain activity. AG may provide sufficient somatosensory and vestibular stimulation to induce neuroplasticity in sensorimotor cortical and cerebellar regions, increasing neural efficiency of these brain regions and facilitating adaptation. These sensorimotor deficits observed during flight may indicate the microgravity interferes with adaptive capacity. We hypothesized that those who received daily AG would maintain better performance during and following HDBR compared with controls. Further, we hypothesized that, compared with controls, the AG group would require lower levels of brain activation in task-related sensorimotor brain regions both during and following HDBR. The measures implemented in this study overlap with those in our ongoing NASA-supported flight and prior bed rest studies (Koppelmans et al., 2013; Yuan et al., 2018a; 2018b; Lee et al., 2019; Hupfeld et al., 2019; 2020; 2021 McGregor et al., 2020; 2021; Salazar et al., 2020; Banker et al., 2021).

## 2. Materials and Methods

### 2.1 Participants and Study Timeline

Twenty-four participants (8 F, 33.3 ± 9.17 yrs, 174.6 ± 8.6 cm, 74.2 ± 10.0 kgs; Table 1), provided their written informed consent to participate in this study and were assigned to one of three groups. Participants were screened through a tolerance test (AG2 protocol; Linnarsson et al., 2015) to ensure they would be able to complete centrifugation. Two groups received centrifugal AG applied either 1) continuously in one 30-minute bout daily (n = 8); or 2) intermittently in six bouts of 5 minutes with 3 minutes between each daily (n = 8). The third group (n = 8) served as a control group (CTRL) that received no AG. There were no statistical differences between the two AG subgroups in either behavior or brain activation; thus, we consider one (n = 16) AG group in all statistical models. All subjects were familiarized with AG twice during the baseline data collection (BDC-11 and BDC-4) prior to being divided into AG and CTRL groups. The University of Florida and NASA Institutional Review Boards as well as the local ethical commission of the regional medical association (Ärztekammer Nordrhein) approved all study procedures.

**Table 1:**
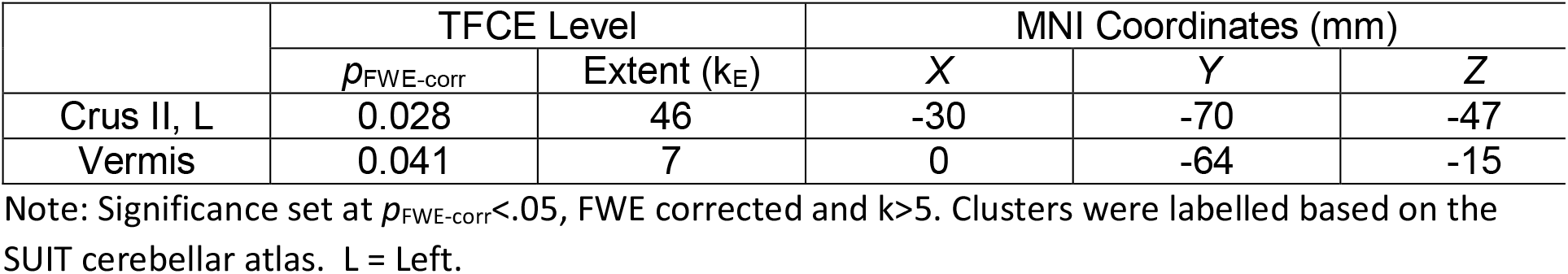
Cerebellar regions that changed during task baseline phase.

Participants completed 60 days of 6° HDBR and performed a range of sensorimotor and cognitive tasks both in and out of the centrifuge at multiple time points pre, during, and post-HDBR (Fig 1.). Both prior and post-HDBR participants were kept on-site for 2 weeks to extensively assess performance, totaling around 88 days of participation.

**Fig 1.**
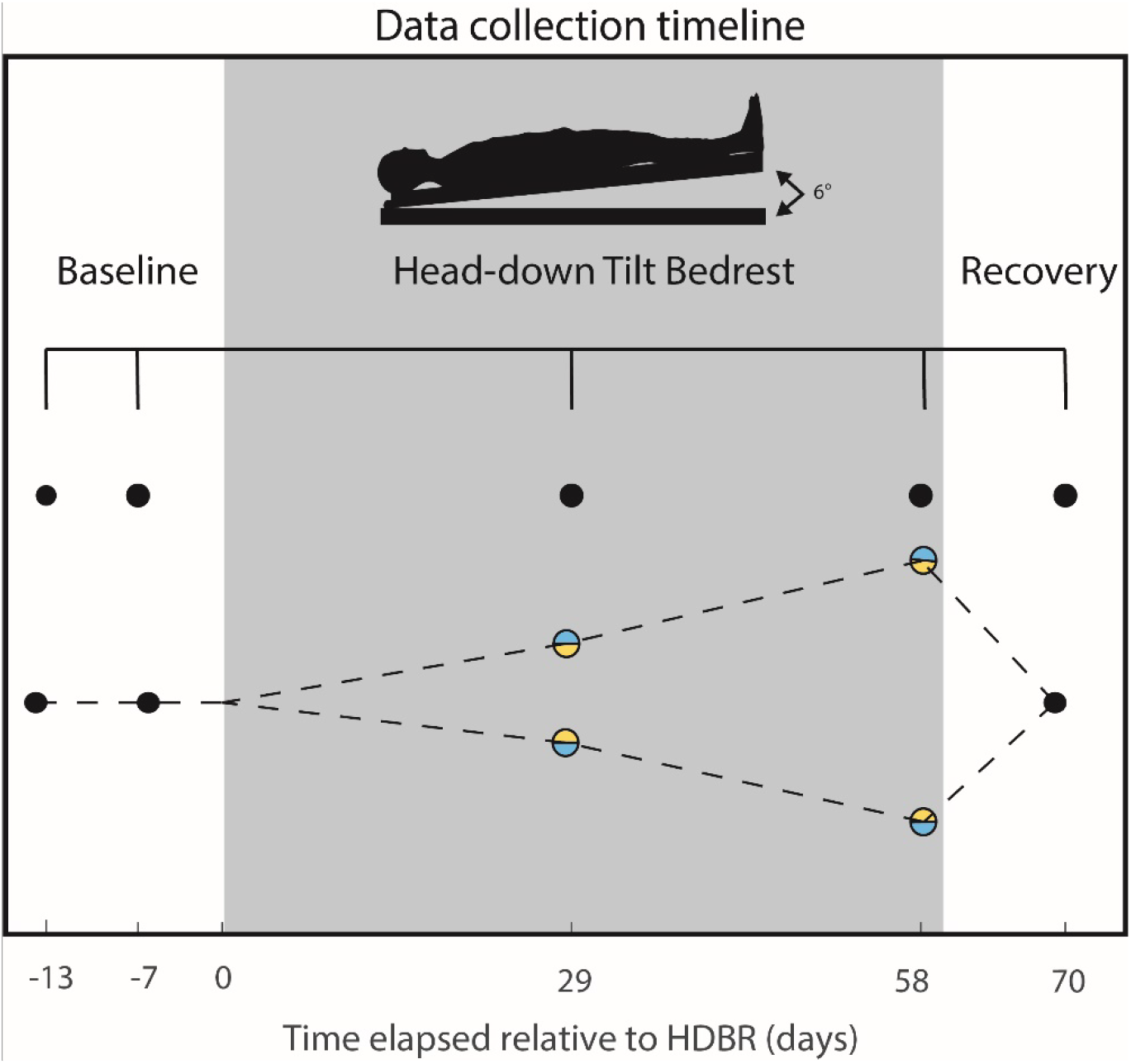
Task timeline. Baseline data was collected twice pre- (-13 and -7 days), twice during (29 and 58 days) and once post-HDBR (+10 days). The dashed line represents our hypothesis regarding brain activation; we hypothesize both groups will have similar patterns of activation prior to HDBR, they will have differential patterns during HDBR due to the presence, or lack thereof, of artificial gravity. Following HDBR, we hypothesized that participants brain activation would return to levels similar as pre-HDBR by 10 days post-HDBR. Black dots represent when we expected both groups to have similar levels of brain activation, and blue/yellow represent where we expected differential brain activation patterns.

### 2.2 Sensorimotor Adaptation Task

During each data collection, participants performed a visuomotor adaptation task inside of a 3T Siemens Magnetic Resonance Imaging (MRI) scanner. Participants moved an MRI-compatible joystick with their right thumb and index finger to reach targets displayed on a screen with real-time feedback of a cursor indicating the joystick location. In the beginning of each trial, a start target would appear in the center of the screen as well as the cursor indicating the joystick location. A target (open circle) would then appear pseudo-randomly in one of four possible locations (above, below, left, or right of start), resulting in an equal number of trials in each target direction. Participants were instructed to move the cursor to the target as quickly as possible and hold the cursor within the target until it disappeared, then to release the joystick and allow the cursor to re-center in the initial starting position.

The task had 4 phases: 1) baseline, 2) early adaptation, 3) late adaptation and 4) re-adaptation. Baseline and re-adaptation consisted of two blocks each (BS1 and BS2, RA1 and RA2, respectively), while early and late adaptation consisted of 4 blocks each (EA1, EA2, EA3 and EA4; LA1, LA2, LA3 and LA4, respectively; Fig. 2). Each block consisted of 16 trials. During the baseline phase, veridical visual feedback was provided for the participants to become accustomed to the basic task, then during adaptation phases a 45° clockwise rotation was applied to the visual feedback. Subjects were not given any explicit instructions or information regarding the rotation; it was introduced to induce an adaptive response for the participants to gradually adapt to across trials. Following that, the rotation was removed during the re-adaptation phase to measure the aftereffects of adaptation. As in our previous work with this paradigm (Ruitenberg et al., 2018; Banker et al., 2021; Salazar et al., 2021), we used direction error as the primary outcome metric. Direction error here is defined as the angle between the start location and the cursor position at the time of peak velocity, and a straight line connecting the start to the target. We also calculated movement time and reaction time for each trial.

**Figure 2.**
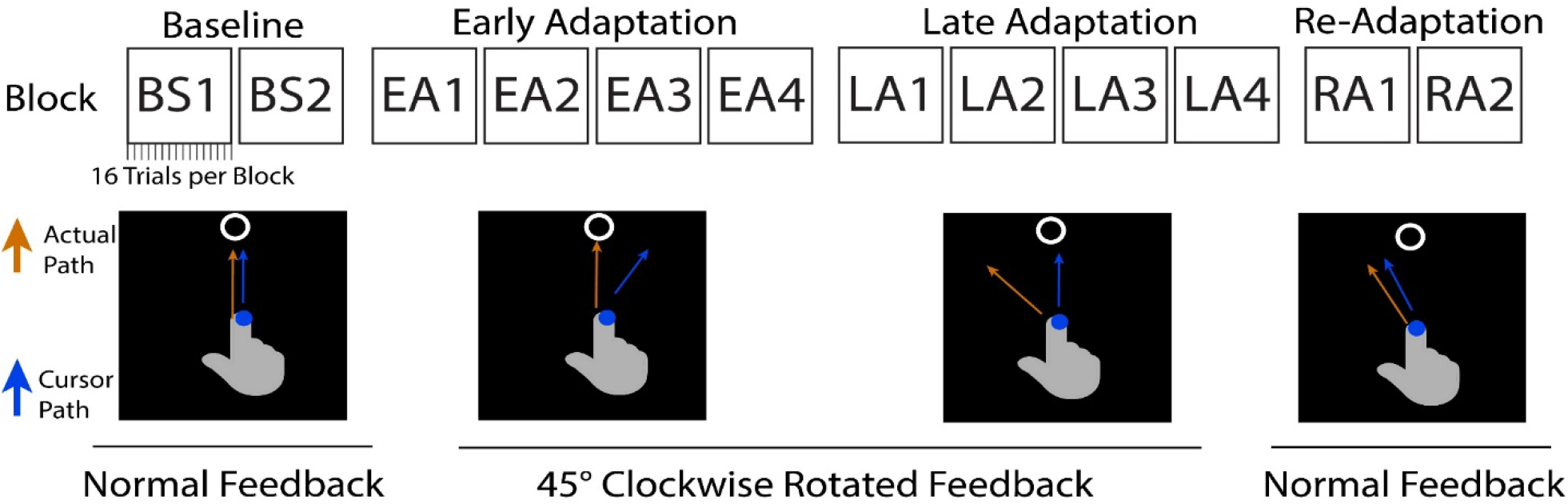
Sensorimotor Adaptation Task. During the two baseline blocks (BS, 16 trials per block) participants move a joystick that controls a cursor on the screen with normal feedback. We induced a 45 degree clockwise rotation of the feedback in the adaptation phases (Early and Late, EA and LA) to create sensory prediction errors that then induce sensorimotor adaptation. Following 8 blocks total, the rotation is removed. Participants then need to re-adapt to normal feedback (RA blocks).

Participants completed 192 trials (16 per block) per session. To see if there were any effects of AG on how fast participants adapted, we calculated learning rates for the early, late and re-adaptation blocks by subtracting the last 4 trials’ average direction error (i.e., last four trials of block LA4) from the first four trials’ average direction error (i.e., first four trials of block LA1) (Taylor et al. 2014; Bond & Taylor, 2015). Additionally, we calculated savings scores by subtracting a later session’s direction error of the first block (EA1) from an earlier session’s direction error of the first block (EA1) (e.g., subtracting BDC-7’s EA1 direction error from HDT29’s EA1). Savings scores were investigated only in the early and re-adaptation phases, as savings is typically not evident in later adaptation trials (Herzfeld et al., 2014; Coltman et al., 2019; Avraham et al., 2021).

### 2.3 fMRI Parameters

All MRI data were acquired with the same 3-Tesla Siemens Biograph MRI scanner located at the :envihab facility. First, we acquired a T1-weighted MPRAGE gradient-echo pulse sequence with the following parameters: TR: 1.9 s, TE: 2.4 ms, flip angle: 9°, FOV: 250 × 250 mm, matrix: 512 × 512, slice thickness: 1.0 mm, voxel size: 0.49 × 0.49 × 1.0 mm, 192 slices. We then acquired fMRI using a gradient echo T2*-weighted echo-planar imaging sequence with the following parameters: TR: 2500 ms, TE: 32 ms, flip angle: 90°, FOV: 192 × 192 mm, matrix: 64 × 64, slice thickness: 3.5 mm, voxel size: 3 × 3 × 3.5 mm, 37 slices. During the fMRI collection, the subject’s body remained in the HDBR position laying upon a foam wedge, however the head was flat and supine within the MRI head coil. The visuomotor adaptation task was performed in a block design, alternating between task trials (40s) and rest (20s) across 4 runs (1 for each BS, EA, LA and RA phase).

### 2.4 Whole Brain fMRI Pre-processing

Pre-processing of the fMRI data were conducted using Statistical Parametric Mapping 12 (SPM12; version 7219; Penny et al., 2011), MATLAB R2019a and Advanced Normalization Tools (ANTs; Avants et al., 2011) and FSL (Jenkinson et al., 2012). This pre-processing procedure is the same as we have used in our past HDBR work (Hupfeld et al., 2019; Salazar et al., 2020; Salazar et al., 2021). First, field maps were created to map and correct for B0 inhomogeneities utilizing the FSL topup tool (Jenkinson et al., 2012). Next, we corrected the images for slice timing and realigned and resliced individual volumes to correct for volume-to-volume head motion in SPM12. We used the Artifact Detection Tool (ART; https://www.nitrc.org/projects/artifact_detect/) to identify volumes with framewise displacement motion ≥ 2.0 mm and global brain signal Z threshold ≥ 9; we covaried out any outlier volumes (see below). We then used ANTs (Avants et al., 2011) to normalize the whole brain images to MNI space in multiple steps. First, we used the ANTs AntsMultivariateTemplateConstruction.sh function to create a longitudinal T1 template for each participant using their five T1-weighted skull stripped structural images (collected at each of the 5 time points). This multistep procedure has proved to normalize the images into a standard space, particularly when accounting for the physical brain changes that occur in HDBR (upward shift of the brain, ventricular expansion, etc.). Second, we created participant-specific mean fMRI templates using the same ANTs function. For each participant, we calculated the mean volume of each preprocessed run, and inputted them into the ANTs AntsMultivariateTemplateConstruction.sh function to create a fMRI template. We then coregistered each subject’s fMRI template and their respective T1 template using the AntsRegistration.sh function. Third, we normalized each participant’s T1 template to MNI152 space using the AntsRegistrations.sh function. Transformations resulting from the above registrations were concatenated into a flow field, and were applied to the native space T1 or preprocessed functional run using ANTs’ ApplyTransforms.sh function. Ultimately, this resulted in fMRI images from each participant and each time point normalized to standard (MNI) space. Finally, we used SPM12 to spatially smooth with an 8 mm full-width at half-maximum three-dimensional Guassian kernel.

### 2.5 Cerebellar Pre-processing

Similar to our previous spaceflight work (Salazar et al., 2020; 2022; Hupfeld et al., 2021; McGregor et al., 2021), we used specialized pre-processing methods for the cerebellum which leveraged the CEREbellum Segmentation (CERES; Romero et al., 2017) pipeline and the Spatially Unbiased Infratentorial Template (SUIT; Diedrichsen, 2006; Diedrichsen et al., 2009). First, we used the CERES pipeline to segment the cerebellum from the rest of the brain in the participant-specific T1 templates. Then, we created a binary cerebellar mask from the CERES native space output using ImCalc in SPM12 and used this to isolate the cerebellum from the participant-specific T1 template with FSLmaths. Then we used ANTs AntsRegistration.sh to transform the T1 cerebellar template to SUIT space. We then transformed the slice timed, realigned and resliced functional runs (described above) into the participants’ T1 template space using AntsTransform.sh and masked with the binary cerebellar mask in FSL. Finally, we normalized the masked, functional runs into SUIT space using AntsApplyTransforms.sh. We then applied a 2 mm full-width half-maximum three-dimensional smoothing Gaussian kernel to SUIT space cerebellar images in SPM12. We chose the 2 mm kernel based on the small lobule size in the cerebellum; this is compatible with other studies (Diedrichsen, 2006; Diedrichsen et al., 2009; 2011).

### 2.6 Sensorimotor Adaptation Behavior Statistical Analyses

We used the nlme package (Pinheiro et al., 2020) in R 3.6.1 (R Core Team, 2019) to fit linear mixed effects models with restricted maximum likelihood (REML) estimation to assess sensorimotor adaptation direction error, movement time and reaction time performance changes over time. In each model we fit a random intercept for each subject to allow for different starting points for each individual. We fit two models: 1) to evaluate the effects of the HDBR and HDBR+AG environment on performance and 2) to evaluate recovery after exiting the HDBR and HDBR+AG environment.

### 2.7 Subject-Level fMRI Statistics

Subject-level brain activity for the visuomotor adaptation task was calculated separately for the whole brain and the cerebellum. For both the whole brain and cerebellum, four statistical maps were created for every participant and time point on a voxel-by-voxel basis according to the following contrasts: 1) baseline > rest, 2) early adaptation > baseline, 3) late adaptation > baseline and 4) re-adaptation > baseline. Initial statistical analysis showed a group difference (discussed below, Section 3.2) in baseline versus rest, thus we decided to investigate early, late and re-adaptation relative to baseline instead of relative to rest. Based on our previous work (Hupfeld et al., 2019; Salazar et al., 2020; 2021), we used a first level masking threshold of - Infinity and masked out non-brain areas using the SPM intracranial volume mask. This allowed us to include all voxels in the subject-level general linear models, as opposed to the SPM default which includes only voxels with a mean value of ≥80% of the global signal.

### 2.8 Group-Level fMRI Statistical Analyses

To assess brain activation changes throughout HDBR+AG we tested multiple group-level statistical models, described in detail below (Sections 2.8.1-2.8.2). Each group-level model was built using the Sandwich Estimator Toolbox (SwE; Guillaume et al., 2014). The SwE toolbox uses a noniterative marginal model to prevent within-subject convergence problems inherent to longitudinal designs, providing optimal analysis of longitudinal MRI data, especially with small data sets and missing data. No modification to the SWE default setup was made other than using non-parametric wild bootstrapping with 999 permutations (Guillaume & Nichols, 2015) and threshold free cluster enhancement (TFCE, Smith & Nichols, 2015). TFCE does not require an arbitrary cluster threshold and instead provides greater sensitivity and stability through an algorithmic boosting of signals without changing the location of their maxima (Smith & Nichols, 2009).

#### 2.8.1. Time Course of Neural Visuomotor Adaptation Response to HDBR+AG

To test the potentially mitigating effect of AG on HDBR-induced changes, we implemented an *a priori* hypothesized weighted longitudinal model (Fig. 1). We created longitudinal contrasts that include all five data collection sessions, with each weighted to test for our hypothesized changes. The models used here tested for linear increases or decreases in brain activity across the HDBR phase that returned to baseline values following bed rest (Fig.1), contrasting the two groups against each other. In other words, we tested to see if there was linearly increasing activation in the CTRL group and linearly decreasing activation in the AG group (and vice versa), identifying regions that they differed in their activation. We have used similar approaches in other HDBR studies (Yuan et al., 2016; 2018a; 2018b; Hupfeld et al., 2019; Salazar et al., 2020; 2021; McGregor et al., 2021) with. Mean centered age and sex were included as covariates. Significance was analyzed at *p*<0.05, family-wise error (FWE) corrected for multiple comparisons. For whole brain analysis, an explicit mask was used to investigate only gray matter effects in the cerebrum. This mask was created through binarizing the Computational Anatomy Toolbox 12 (CAT12; Dahnke et al., 2013; Gaser & Kurth, 2017) MNI-space gray matter template at a threshold of 0.1 and masking out the cerebellum using the SUIT cerebellar template. Cerebellar analyses were conducted only on the cerebellum as discussed above in “Cerebellar Pre-processing” with no mask utilized for analysis.

#### 2.8.2. Brain-Behavioral Correlation of Visuomotor Adaptation Response to HDBR+AG

To assess the effects of AG on HDBR we used a similar approach in brain-behavioral correlations as discussed above. However, here we assessed the change in direction error between BDC-7 to HDT58 and the changes in brain activation at the same time points and contrast the groups against each other. Results masks from the longitudinal analysis were also utilized here to limit the search space.

## 3. Results

### 3.1. Sensorimotor Adaptation Behavior Results

Direction error (DE) showed no changes in response to HDBR or HDBR+AG during the baseline (*p*=0.28), early (*p*=0.89), late (*p*=0.23) or re-adaptation (*p*=0.42) phases of the sensorimotor adaptation task (Supplementary Table 1). A main effect of trial was identified for early adaptation (*p*<0.0001), late adaptation (*p*<0.0001) and re-adaptation (*p*<0.0001) indicating that DE decreased as participants responded to the perturbation of the task. There was also a main effect of block during early (*p*=0.0002) and late (*p*<0.0001) adaptation, reflecting improvements with practice. DE for the two groups across all 5 time points is displayed in Figure 3.

**Figure 3.**
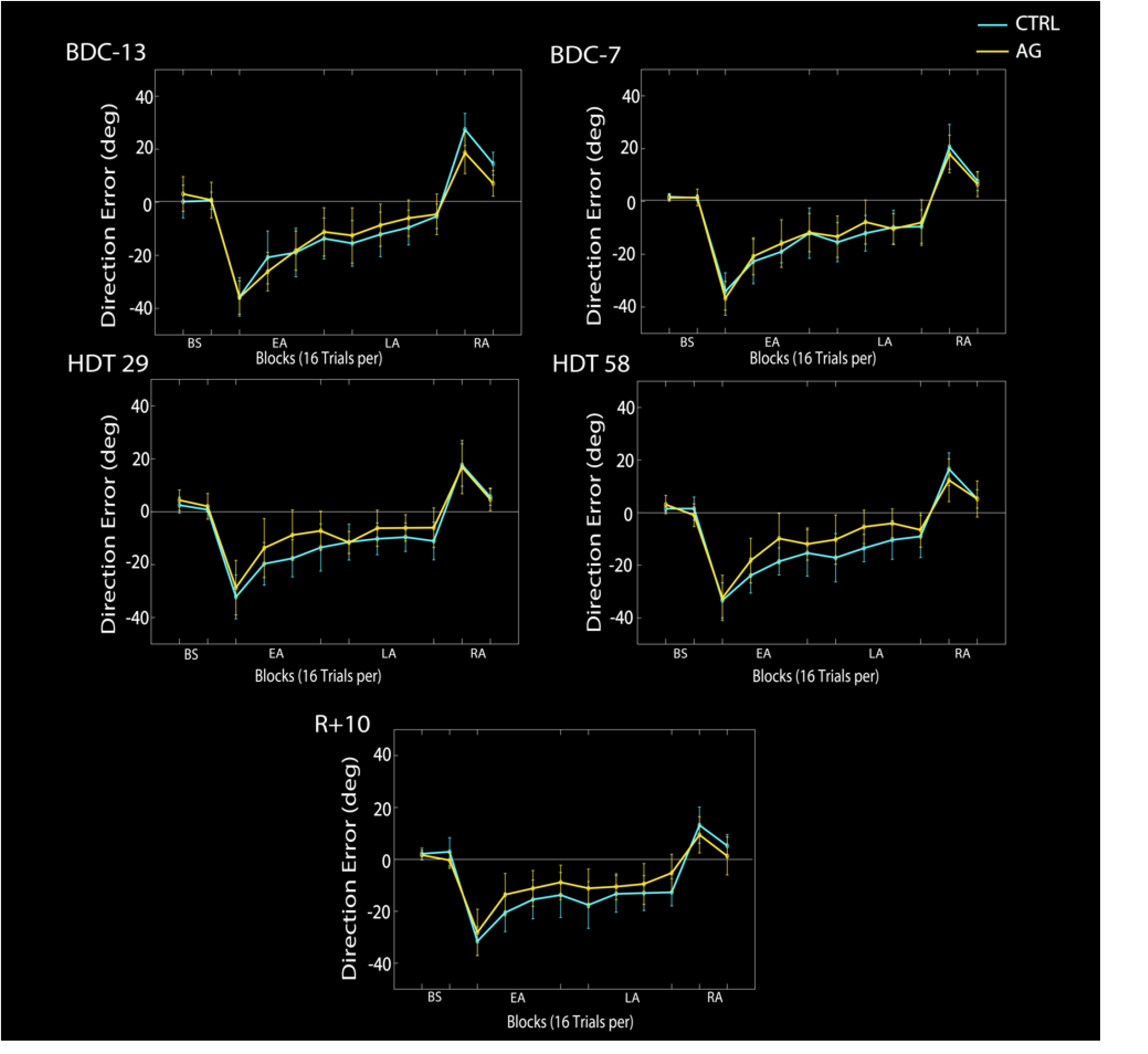
Direction error (DE) performance at all five time points for both the AG (yellow) and CTRL (blue) groups. Although there was a visual trend for the AG group to perform better after entering HDBR (and starting AG), DE was not significantly different between the two groups over time. BDC-13 and BDC-7 are 13 and 7 days prior to entering in HDBR. HDT 29 and HDT 58 are 29 and 58 days in HDBR while receiving daily artificial gravity in the AG group. R+10 is 10 days following the conclusion of HDBR. Values are mean across subjects with error bars illustrating standard error. This task was performed inside an MRI scanner.

There were no significant group by time interactions for reaction time in baseline (*p*=0.08), late adaptation (*p*=0.50) or re-adaptation (*p*=0.98). During early adaptation, there was a significant three-way interaction between group, time spent in HDBR, and block for reaction time (*p*=0.05). That is, during HDBR those that received AG had quicker reaction times initially, and these reaction times got faster over HDBR. Further, there were main effects of trial for each phase (*p*<.0001), as participants reacted faster with practice. During baseline, there was also a main effect of sex (*p*=.04), where men reacted more quickly. There were additional main effects of block during early (*p*<0.0001) and late adaptation (*p*<0.0001), as participants also reacted more quickly with practice across blocks.

There were no significant group by HDBR interactions for movement time in any of the four task phases (*p*=0.69; *p*=0.85; *p*=0.79; *p=*0.26). There was a main effect of sex in the baseline phase, in which men moved at a quicker speed (*p*=0.03), in addition to a main effect of age in the early adaptation phase (*p*=0.02), in which older individuals moved slower. Further, we observed main effects of trial in all four phases (all *p*<0.0001), and block for late adaptation (*p*=0.03), reflecting faster movement times with practice.

We also investigated whether there may be a difference in learning rates and savings between the AG and CTRL groups. Savings is defined as improved adaptation to a perturbation when you have previously experienced it (Ruitenberg et al., 2018) and has been shown to last from one day after initial learning (Bédard et al., 2011; Seidler et al., 2017; Villalta et al., 2015) up to one year (Landi et al., 2011; Yamamoto et al., 2006). Similar to what we observed with the trial by trial analyses, there were no significant differences in learning rates in all four phases (*p*=0.19; *p*=0.60; *p*=0.61; *p*=0.80) between groups or across the data collection sessions. There were also no significant differences in savings scores for early adaptation (*p*=0.55) or re-adaptation (*p*=0.43), suggesting that savings did not occur in either group across HDBR, nor from BDC-13 to BDC-7.

### 3.2. HDBR+AG Neural Correlates of Visuomotor Adaptation

In the baseline phase of the task, we identified two clusters in the cerebellum that significantly differed in their activity from pre-to in- and post-bed rest between groups (*p*_FWE-corr_<0.05, cluster size k>5; Table 1). In these clusters, we found that the CTRL group showed an increase in cerebellar activation during HDBR that resolved by 10 days post-HDBR (Fig. 4). Those in the AG group showed a decreasing or stable pattern of activation throughout HDBR+AG that did not change following the conclusion of HDBR. There were no significant differences between the groups in the other three phases relative to rest.

**Figure 4.**
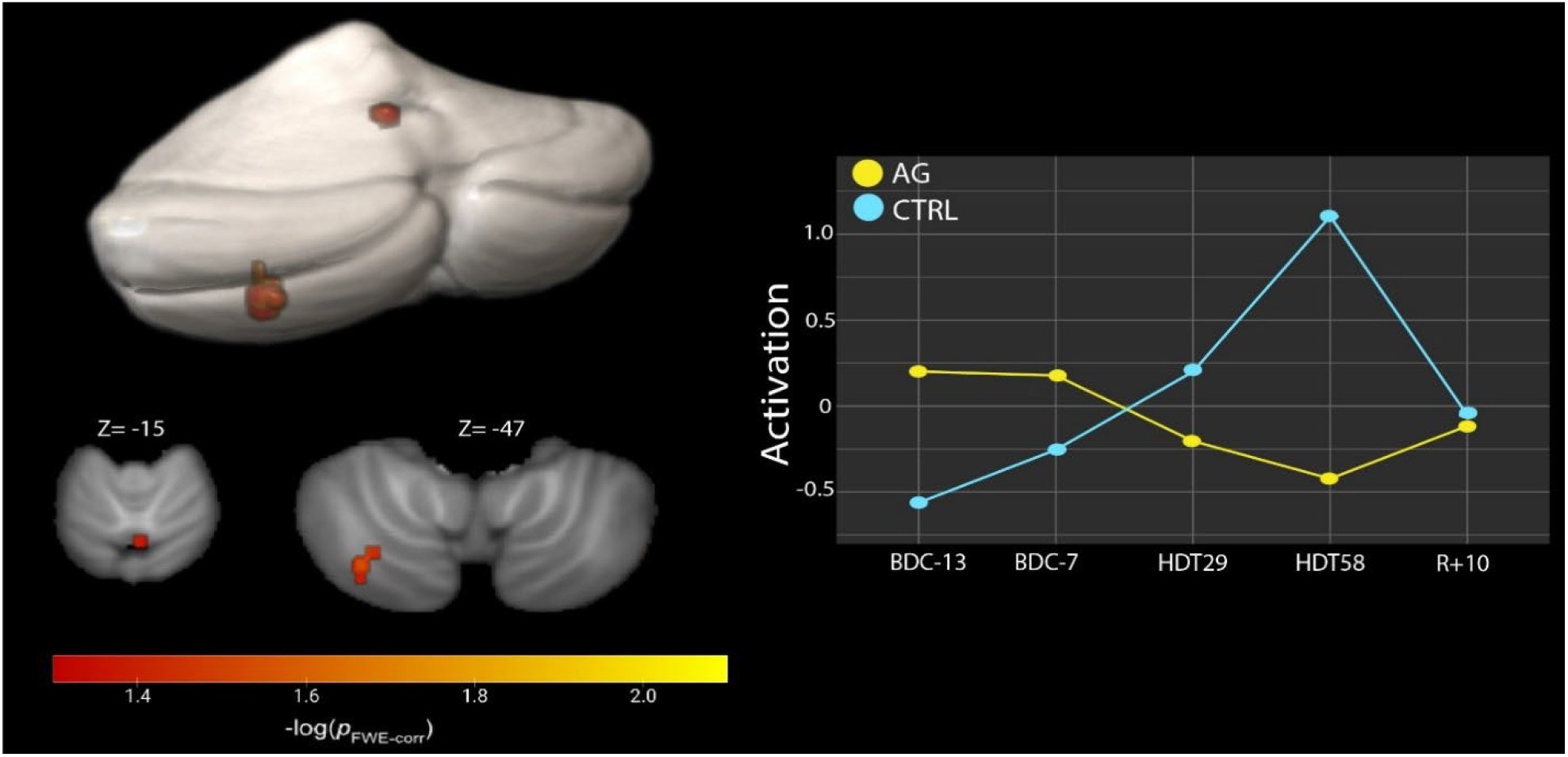
Group differences in activation within the cerebellum during the baseline phase (normal visual feedback) of the adaptation task. The AG group showed a differential response of decreasing activation during the task when they first entered into the HDBR+AG environment. In contrast, CTRL participants showed increasing activation throughout HDBR that returned toward baseline levels by 10 days post-HDBR. The activation profile for the L Crus II is shown above, right, the vermis profile (not shown) is similar. Results are overlaid onto the SUIT cerebellar template. *p*_FWE-corr_, brighter colors represent smaller p-values.

When analyzing brain activation in relation to baseline, we found six clusters that differed significantly between the two groups in the early adaptation phase (*p*_FWE-corr_<0.05, cluster size k>10; Table 2; Fig 5.). Within these clusters we observed that, specific to HDBR, those in the CTRL group overall increased their activation relative to baseline. In contrast, the AG group showed decreasing activation in these regions throughout HDBR, which then returned to baseline level 10 days after ending bed rest.

**Table 2:** Brain regions that displayed longitudinal changes in early activation that differed between groups.

|  | TFCE Level |  | MNI Coordinates (mm) |  |  |
| --- | --- | --- | --- | --- | --- |
| | $p_{\text{FWE-corr}}$ | Extent ($k_E$) | X | Y | Z |
| L Calcarine | 0.031 | 576 | 2 | -86 | 10 |
| L Cuneus | 0.034 | 395 | -8 | -102 | 4 |
| L Thalamus | 0.035 | 75 | -22 | -20 | 8 |
| L Superior Frontal Gyrus / preSMA | 0.044 | 135 | -14 | 16 | 54 |
| L SMA | 0.047 | 57 | -12 | 4 | 60 |
| R Cuneus | 0.048 | 13 | 4 | -84 | 30 |

Note: Brain regions that showed longitudinal changes between AG and CTRLs. Significance set at $p_{\text{FWE-corr}} < 0.05$ and $k > 10$ . Clusters were labelled using the AAL3 atlas. Clusters are listed from smallest $p$ value to largest. L = Left, R = Right.

**Figure 5.**
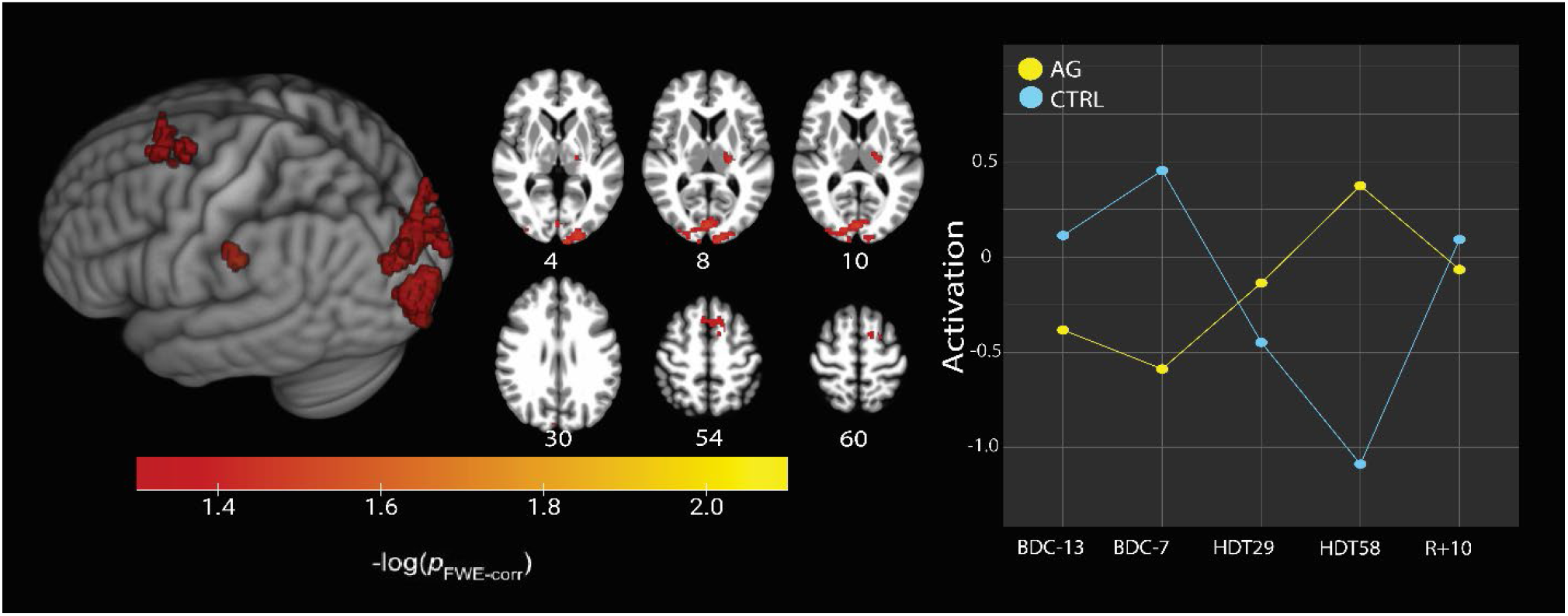
Activation differences between the two groups during early adaptation relative to baseline. The activation profile for the L Cuneus region is shown on the right, which is similar to activation profiles of other significant clusters. Results are overlaid onto the MNI standard space template. *P*<.05, FWE corrected, brighter colors represent smaller p-values.

### 3.5 Brain-Behavioral Correlation of Visuomotor Adaptation Response to HDBR+AG

We further tested for associations between performance and brain activation within the cerebellar regions listed in Table 1 that displayed brain activation changes during the baseline portion of the task. We identified two clusters that displayed a significant time by group interaction in longitudinal changes from BDC-7 to HDT58 that correlated with direction error changes over this time (TFCE<0.05; Table 3). That is, subjects in the CTRL group showed little change in brain activation regardless of change in performance. However, within the AG group those subjects who improved their performance during baseline (normal visual feedback) trials from pre-HDBR to late-HDBR exhibited a decrease in brain activation while performing the task that was correlated with their performance changes.

**Table 3:**
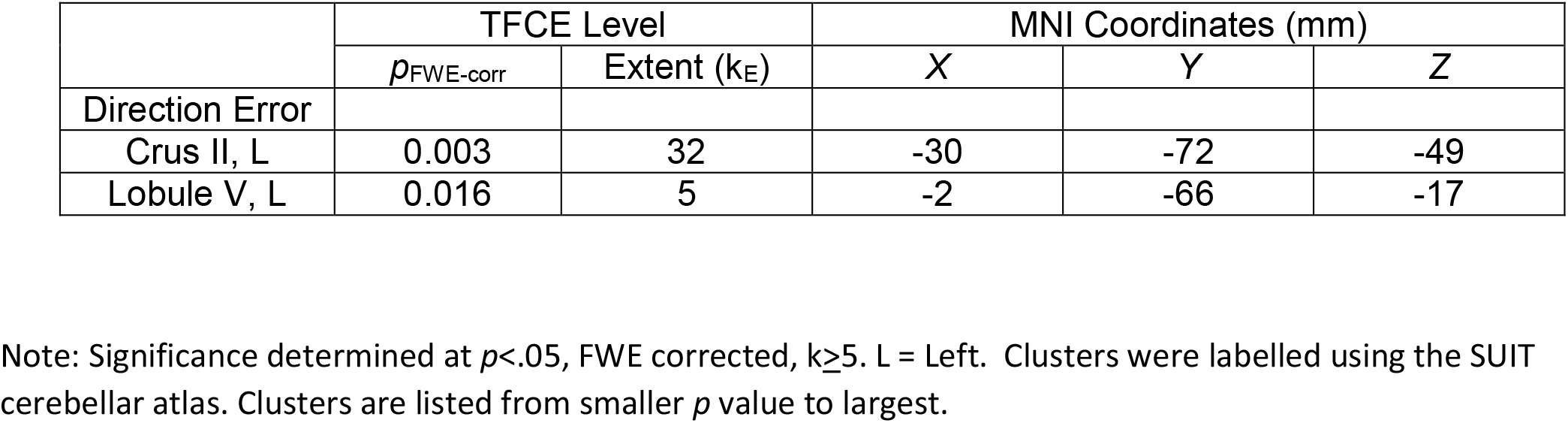
Cerebellar brain and baseline phase behavior correlation

|  | TFCE Level |  | MNI Coordinates (mm) |  |  |
| --- | --- | --- | --- | --- | --- |
| | $p_{FWE-corr}$ | Extent ($k_E$) | X | Y | Z |
| Direction Error |  |  |  |  |  |
| Crus II, L | 0.003 | 32 | -30 | -72 | -49 |
| Lobule V, L | 0.016 | 5 | -2 | -66 | -17 |

Note: Significance determined at $p < .05$ , FWE corrected, $k \geq 5$ . L = Left. Clusters were labelled using the SUIT cerebellar atlas. Clusters are listed from smaller $p$ value to largest.

Within the clusters showing differential group changes over time for the early adaptation phase of the task, we identified a brain and behavior correlation that differed for the AG versus CTRL groups. Specifically, direction error during early adaptation and activation levels in the right calcarine and left cuneus (Table 4) were associated from pre-to late-HDBR. Those in the AG group overall performed better (i.e., lower direction error) at late-HDBR and had no association between brain activity changes and DE changes (Fig. 6). CTRLs that had increased activation at late-HDBR had worse performance (i.e., higher direction error) at the same time. CTRLs that performed more similarly to those that received AG, had lower brain activation and better performance.

**Table 4:** Brain and behavioral correlation during early adaptation within regions that showed longitudinal change.

|  | TFCE Level |  | MNI Coordinates (mm) |  |  |
| --- | --- | --- | --- | --- | --- |
| | $p_{FWE-corr}$ | Extent ($k_E$) | X | Y | Z |
| R Calcarine | 0.006 | 523 | 14 | -88 | 10 |
| L Cuneus | 0.038 | 77 | -8 | -98 | 8 |

Note: Significance determined at $p_{FWE-corr}$ , $k>10$ . L = Left, R = Right. Clusters were labelled using the AAL3 atlas. Clusters are listed from smaller $p$ value to largest.

**Figure 6.**
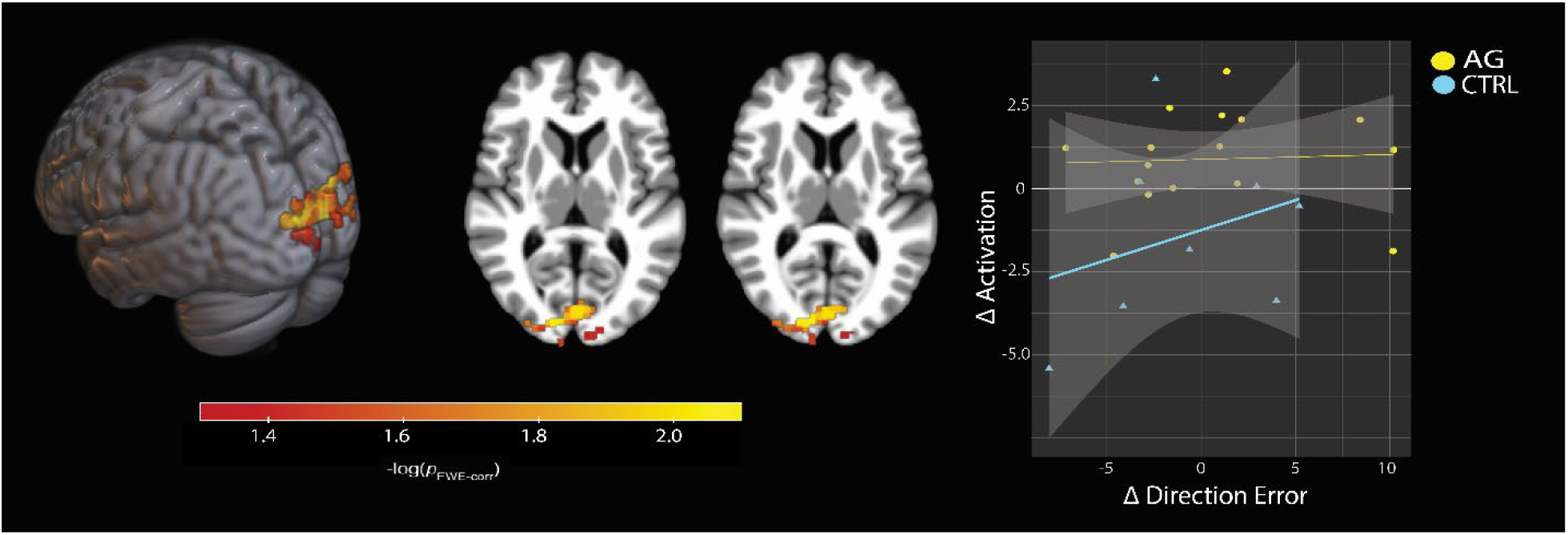
Brain and behavior correlation of direction error during the EA phase and functional brain activity during EA. Positive change in direction error indicates improvement in performance (i.e., decreasing error), where positive change in activation indicates decreased activation by late-HDBR. Overall, CTRL subjects that increase their behavioral performance by late-HDBR have decreased brain activation, performing more similarly to AG participants.

## 4. Discussion

This study investigated the efficacy of AG as a countermeasure to mitigate HDBR-induced deficits in sensorimotor adaptation. We observed no statistical differences in sensorimotor adaptation between those that received AG and those that did not. However, we did identify cerebellar activation differences between the AG and CTRL groups during the baseline phase of the task (i.e. when performing the task with no feedback rotation); this group difference started at the onset of HDBR, suggesting that AG may influence the neural control of basic motor processes. Further, we identified differential responses between AG and CTRL participants during the early adaptation phase within various sensory processing and integration regions. Those who received AG had decreased functional brain activation whereas CTRLs showed increases in brain activation in these regions. These changes were correlated with behavioral performance such that CTRLs who had reduced brain activation by late-HDBR, similar to the AG group, had increased performance. Taken together, these findings suggest that AG may increase neural efficiency during manual sensorimotor adaptation tasks. Thus, AG should be further explored as an integrated countermeasure for mitigating spaceflight induced brain changes.

We identified a significant difference in cerebellar brain activation during the baseline portion of the task, specific to when participants were in HDBR. That is, the AG group showed decreased activation as compared to CTRLs who had increased activation that returned to baseline level post-HDBR. Interestingly, the AG participants with the lowest levels of cerebellar involvement were the most accurate at the task under baseline conditions. Portions of the cerebellum play a role in error processing, which may explain this brain-behavior association (Kitazawa et al., 1998; Seidler et al., 2004; Diedrichsen, 2005). There has also been evidence that Crus II is structureally connected to the parietal lobe, which has been known to play a major role in sensory integration (Clower et al., 1996; Sasaki et al. 1975). White matter tract-tracing studies in non-human primates have also demonstrated structural connectivity between the posterior parietal cortices and Crus I and II (Hoover & Strick, 1999; Kelly & Strick, 2003).

Further, the motor cortex has been shown to project to the vermis and play a role in postural adjustment (Coffman et al., 2011). The vermis has also been identified to functionally change in spaceflight, and has been thought to process graviceptive inputs (Yates et al., 2003; Krasnov & Krasnikov, 2009). Functionally, cerebellar lesions have been shown to result in difficulty with reaching (Glickstein et al., 2005) and impair smooth pursuit eye movement (Westheimer & Blair; 1973), a key aspect of sensorimotor adaptation. Here, during the baseline phase, we found differential activation in the vermis and Crus II, both regions that are associated with sensorimotor behavior. AG has been theorized to serve as a countermeasure for the lack of gravity by providing sensory inputs that are absent during microgravity. Thus, it is possible that the decreased cerebellar activation during baseline is the result of increased sensory inputs from artificial gravity, replacing those that are typically absent during HDBR, and leading to neuroplastic changes that do not occur in the CTRL group.

We also identified differential activation responses between groups in the left hippocampus, left supplementary motor area, left superior frontal gyrus (primarily overlapping with the pre supplementary motor area), left calcarine and left & right cuneus. In these regions, the AG group had decreased activation relative to baseline during the HDBR+AG phase, and the CTRLs had increased activation relative to baseline. These regions are known to be related to adaptation, voluntary movement, sensory integration and higher order cognition. The supplementary motor area has been shown to contribute to planning and execution of voluntary movement (Picard & Strick, 2003; c.f. Nachev et al., 2008). The hippocampus has been argued to serve a sensorimotor-gating mechanism (Bast & Feldon, 2003) and is known to receive sensory information, particularly vestibular projections (Smith, 1997; Hitier et al., 2014) and process stimuli during sensorimotor tasks (c.f. Crochet et al., 2019). The cuneus is well known to play a role in visual processing tasks (Malach et al., 1995), and has been shown to have increased connectivity with other task related regions in professions that require high visuomotor and visuospatial skills, such as gymnasts (Huang et al. 2015) and commercial airline pilots (Qiu et al., 2021). Additionally, the pre-SMA region has been shown to be play a role in motor planning (Shimzu et al., 2020), as well as with processing and movement execution (Hoshi & Tanji, 2004; Cona et al., 2016). Further, the pre-SMA plays a role in the temporal and spatial processing of voluntary movement (Mita et al., 2009; Kotz & Schwarte, 2011). These regions showed HDBR specific increases in brain activation in the CTRL participants while performing the early trials of the sensorimotor adaptation task, however in the AG group they show decreased activation. This may be due to increased sensory information to these and connected regions from the AG application. Any resulting sensory / sensorimotor plasticity occurring from the daily AG may then facilitate or interact with neural control during performance of the sensorimotor adaptation task.

While there were no behavioral performance differences between the AG and CTRL groups there was decreased activation relative to the baseline portion of the task (aiming under normal visual feedback) in the AG group. That is, the CTRL group recruited additional neural resources compared to controls to maintain the same level of performance. Within these regions, we identified a brain-behavior correlation that differed between groups. Those that received AG exhibited a decrease in sensorimotor brain activity by late HDBR, and increased performance. Additionally, CTRLs that had similar brain activation changes as seen in the AG group, performed more closely to their level. We believe these findings to indicate that AG may reduce neural cost of sensorimotor performance and increase neural efficiency. This suggests that AG may mitigate the increased neural load of HDBR and increase neural efficiency. Neural efficiency has been investigated previously in other contexts; for instance Jaeggi et al., (2007) found that higher performing participants and lower performing participants (differentiated by a median split) had similar behavioral performance on a cognitive n-back task, yet the high performers had significantly lower brain activation in task-related regions, particularly when the difficulty of the task increased. In skilled athletes, similar results were found compared to novices, with decreased cortical activation in task-related regions during a social cognition task for the experts (Del Percio et al., 2007). Further, it has been shown that skilled athletes had lower brain activation than non-athletes overall on a visuo-spatial task within task-related brain regions (Guo et al., 2017). Here, while we did not identify differences in behavioral performance between groups, the AG group exhibited decreasing brain activity during task performance throughout HDBR. Although speculative, we interpret this as better neural efficiency, particularly in brain regions that may be stimulated by the AG such as the cerebellum.

This work was part of the AGBRESA campaign, a joint analog mission between American, European and German space organizations (NASA, ESA and the DLR) that investigated the effects of AG as an integrated countermeasure across a wide variety of physiological and neurological domains, including the current work. De Martino et al. (2021) analyzed the effects of AG on upright balance, finding that intermittent AG partially mitigated balance impairment from pre to post-HDBR compared to CTRLs, but continuous AG did not differ from CTRLs. This finding is somewhat in line with our own previous findings that AG may be beneficial for balance (Tays et al., 2022). We have also recently reported effects of AG on cognitive function, showing that during centrifugation there was an increased level of cognitive ability compared to controls, as measured by the Paced Auditory Serial Addition Task (PASAT; Tays et al., 2022). Basner et al., (2022) found that HDBR overall had a negative effect on cognitive performance, and that receiving AG did not mitigate these declines. However, the PASAT was administered only during active centrifugation, whereas the other tasks were measured across the HDBR period. The cognitive effects of HDBR and spaceflight have been difficult to identify quantitatively, but it has been suggested that they are due to a lack of neural resources from sensory re-weighting and adapting to the environmental change. Thus leaving a shortage for complex cognitive function. Overall, it is possible that one of the primary benefits of AG is increased sensory input, and thus the ability to counteract HDBR induced sensorimotor declines and some cognitive processes, particularly during active centrifugation. However, further work is needed to understand the mechanistic effects of AG on human physiology and to determine the ideal dosing.

The sample size of this study is limited, making it difficult to identify more subtle group differences, particularly between the pooled AG subgroups. HDBR studies are relatively short compared to the much longer time astronauts spend in microgravity. This may result in only partial adaptive dysfunction compared to astronauts completing International Space Station missions, which typically last about 6 months. We are currently studying this same visuomotor adaptation task and associated brain activity in a group of astronauts (Koppelmans et al., 2013) and thus will be able to assess whether HDBR is an appropriate analog for this task. It is also possible that 30 minutes of AG daily is not sufficient to fully counteract deficits induced by HDBR and / or spaceflight.

Here, we investigated the efficacy of AG as an integrated countermeasure to declines that occur in a spaceflight analog environment (60 days HDBR). The group that received AG had decreased cerebellar activation during HDBR in the baseline portion of the task in contrast to controls, who increased cerebellar activity during the same time. Additionally, we identified that the AG group had decreased activation relative to baseline in the early adaptation phase of the task, whereas controls increased cerebellar activity; despite these differences in brain activity, there were no group differences in adaptation performance. These findings suggest that the sensory stimulation that occurs during AG may result in cortical sensory plasticity, leading to enhanced neural efficiency. These promising findings support further investigation into AG as an integrated countermeasure for the microgravity environment.

## Acknowledgements

This work was supported by grants from the National Aeronautics and Space Administration (NASA 80NSSC18K0783) to R.S., A.M., S.W., and J.B. During the completion of this work G.T. was supported by the University of Florida’s (UF) Graduate Student Funding Award and by NIH T32-NS082128. KH was supported by a National Institute on Aging fellowship 1F99AG068440. H.R.M. was supported by a Natural Sciences and Engineering Research Council of Canada (NSERC) Postdoctoral Fellowship and a NASA Human Research Program Augmentation Grant. The authors would like to thank all of the participants who volunteered their time, without them this project would not have been possible.

